# Pooling nasopharyngeal swab specimens to increase testing capacity for SARS-CoV-2

**DOI:** 10.1101/2020.05.22.110932

**Authors:** Cole Anderson, Fritz Castillo, Michael Koenig, Jim Managbanag

**Affiliations:** Department of Pathology, Landstuhl Regional Medical Center, United States Army

**Author notes:** Address correspondence to Cole Anderson.

## Abstract

The recent emergence of SARS-CoV-2 has lead to a global pandemic of unprecedented proportions. Current diagnosis of COVID-19 relies on the detection of SARS-CoV-2 RNA by RT-PCR in upper and lower respiratory specimens. While sensitive and specific, these RT-PCR assays require considerable supplies and reagents, which are often limited during global pandemics and surge testing. Here, we show that a nasopharyngeal swab pooling strategy can detect a single positive sample in pools of up to 10 samples without sacrificing RT-PCR sensitivity and specificity. We also report that this pooling strategy can be applied to rapid, moderate complexity assays, such as the BioFire COVID-19 test. Implementing a pooling strategy can significantly increase laboratory testing capacity while simultaneously reducing turnaround times for rapid identification and isolation of positive COVID-19 cases in high risk populations.

## Introduction

In December 2019, an outbreak of pneumonia with unknown origin began in Wuhan city, the capital of Hubei province in China^1^. The following month, Chinese researchers had isolated a novel coronavirus, severe acute respiratory syndrome coronavirus 2 (SARS-CoV-2), from patients with viral pneumonia^2^. Pneumonia associated with SARS-CoV-2 was later designated as coronavirus disease 2019 (COVID-19) by the World Health Organization in February 2020^3^. It was determined that after a zoonotic transmission event in Wuhan city^4^, widespread person-to-person transmission quickly occurred that led to the infection and death of over 80,000 and 3,000 people in China, respectively. To date, according to the WHO, there have been 4,258,666 reported cases of COVID-19, including 294,190 deaths worldwide^5^.

Since the initial outbreak in China, COVID-19 has been declared a global pandemic affecting at least 216 other countries, territories or areas. To monitor and diagnose COVID-19, the US Food and Drug Administration (FDA) approved an emergency use authorization (EUA) for the CDC 2019-nCoV Real-Time RT-PCR Diagnostic Panel on February 4, 2020^6^. This protocol allows for the rapid detection of SARS-CoV-2 RNA from clinical specimens such as, nasopharyngeal and oropharyngeal swabs, sputum, bronchoalveolar lavage, and tracheal aspirates. As evidenced by the ongoing SARS-CoV-2 pandemic, increased demand for testing can overwhelm diagnostic laboratories and lead to drastic shortages in supplies and reagents. A strategy to overcome high testing demand is to pool specimens before RNA extraction, test pools, and then retest individual specimens from positive pools. Similar strategies have shown to increase testing capacity for the detection of common infectious diseases such as influenza, HIV, Hepatitis, and *Chlamydia trachomatis*^7–11^.

In this study, we examined the feasibility of pooling nasopharyngeal swab specimens submitted for COVID-19 testing using the CDC 2019-nCoV RT-PCR diagnostic panel without compromising clinical sensitivity. Our data shows that pooling respiratory samples during times of increased volume and low disease prevalence can save time and reagents without significant modifications to laboratory infrastructure or workflow.

## Methods

This study was determined to meet the exempt criteria listed in 32CFR219.104(d) from the Landstuhl Regional Medical Center Exempt Determination Official.

During an outbreak cluster of SARS-CoV-2 in Stuttgart, Germany, 494 nasopharyngeal (NP) swabs were collected and placed into 1.0 ml of normal saline. Specimens were submitted to the Virology laboratory at Landstuhl Regional Medical Center for routine SARS-CoV-2 testing using the CDC 2019-nCoV RT-PCR assay. Post clinical testing, specimens were de-identified and randomly assigned into pools of 10 to create 50 distinct pools (the 50^th^ pool contained 4 specimens diluted in 0.6 ml of transport media). Pools were created by combining 100 ul of each specimen to create 1.0 ml pools. Viral transport media was added to each pool at a 1:1 ratio for nucleic acid extraction performed on the Roche MagNA Pure 24 platform using the MagNA Pure 24 Total NA Isolation kit (Roche). Elution volume was set to 50 ul to concentrate viral RNA. Each round of extraction contained a human specimen control to monitor for PCR inhibition and specimen quality.

Detection of SARS-CoV-2 was performed using the CDC RT-PCR COVID-19 assay, which contains primers and Taqman probes for two specific regions of the SARS-CoV-2 nucleocapsid (N) gene and the human Rnase-P (RP) gene, which is used as an internal positive control for human nucleic acid. PCR was performed according to the CDC protocol using the TaqPath 1-Step RT-qPCR Master Mix, CG kit (Life Technologies) on the Applied Biosystems (ABI) 7500 Fast realtime PCR system. PCR results were interpreted as recommended in the CDC RT-PCR COVID-19 instructions for use. A pool was considered positive if the CT was less than 40. Detection of SARS-CoV-2 in pooled samples using the BioFire COVID-19 Test was performed according to the manufacturer’s instructions for use on the BioFire FilmArray 2.0 and FilmArray Torch systems. All statistical analyses were conducted using Graphpad Prism 6.0.

## Results

The prevalence for individual clinical samples was 4% (19/494) for SARS-CoV-2 RNA. Among the pooled samples, 30% (15/50) were positive for SARS-CoV-2 RNA, while the remaining 70% (35/50) did not have detectable levels of SARS-CoV-2 RNA (**Table 1**). We observed one inconclusive RT-PCR result in our pooled analysis as defined by amplification of only a single SARS-CoV-2 target. In this case, the N2 target for pool 41 failed to amplify while N1 was detected with a relatively high CT (37.3). There were no invalid reactions in our analysis, as defined by reactions where Rnase-P failed to amplify. Out of the 15 positive pools, 4 pools contained 2 positive specimens, while the remaining 11 pools contained only 1 positive specimen (**Table 2**). The mean C_T_ value and standard deviation for N1 and N2 of the pools were 29.2 (4.4) and 29.4 (4.3), respectively. Similarly, the mean CT values of individual positive specimens were 28.0 (4.5) and 29.9 (4.8) for N1 and N2, respectively. Despite dilution, there was no significant difference in mean CT value between the pooled and individually tested specimens (**Figure 1**).

**Table 1.** RT-PCR C_T_ values of pooled specimens positive for SARS-CoV-2

| Positive Pool | Ct Values |  |  |
| --- | --- | --- | --- |
|  | N1 | N2 | Rnase-P |
| 2 | 28.6 | 29.7 | 31.8 |
| 8 | 25.4 | 26.2 | 25.7 |
| 11 | 35.9 | 38.7 | 33.7 |
| 12 | 29.7 | 31 | 32.5 |
| 13 | 30.7 | 32.1 | 31.4 |
| 15* | 29.5 | 30.3 | 30.1 |
| 18 | 27.1 | 27.6 | 25.2 |
| 19 | 20.7 | 20.9 | 25.1 |
| 20* | 26.3 | 26.8 | 25.2 |
| 22 | 29.5 | 30.3 | 30.1 |
| 25 | 30 | 29.3 | 25.8 |
| 41*† | 37.3 | Und. | 26.5 |
| 43* | 23.5 | 23.5 | 27 |
| 47 | 30.9 | 31.3 | 26.2 |
| 50 | 33.6 | 33.3 | 26.2 |

\* Pool containing 2 positive specimens
† Inconclusive RT-PCR result
Und.: Undetected

**Table 2.** RT-PCR C_T_ values of individual specimens positive for SARS-CoV-2

| Pool | Sample | Ct Values |  |  |
| --- | --- | --- | --- | --- |
|  |  | N1 | N2 | Rnase-P |
| 2 | 2696 | 25.2 | 26.2 | 27.6 |
| 8 | 2610 | 24.7 | 26.1 | 28.6 |
| 11 | 2535 | 31 | 34.7 | 25.9 |
| 12 | 2697 | 23.9 | 34.9 | 30.4 |
| 13 | 2624 | 21.1 | 21.9 | 28.3 |
| 15 | 2595 | 29.3 | 30.6 | 29.5 |
| 15 | 2620 | 26.3 | 31.6 | 29.1 |
| 18 | 2586 | 29.4 | 32.1 | 26.7 |
| 19 | 2803 | 20.4 | 21.3 | 27.9 |
| 20 | 2665 | 30 | 31.5 | 28.6 |
| 20 | 2785 | 27.5 | 29 | 25.6 |
| 25 | 2662 | 29.5 | 30.6 | 27.8 |
| 41 | 3010 | 33.5 | 32.2 | 28 |
| 41 | 2975 | 36.1 | 36 | 27.1 |
| 43 | 3186 | 35 | 38.4 | 29.3 |
| 43 | 3202 | 24.2 | 25.3 | 32.1 |
| 47 | 3164 | 30.2 | 31.8 | 30 |
| 50 | 3104 | 31.3 | 30.5 | 22.9 |
| 22 | 2497 | 23.2 | 23.4 | 26.4 |

**Figure 1.**
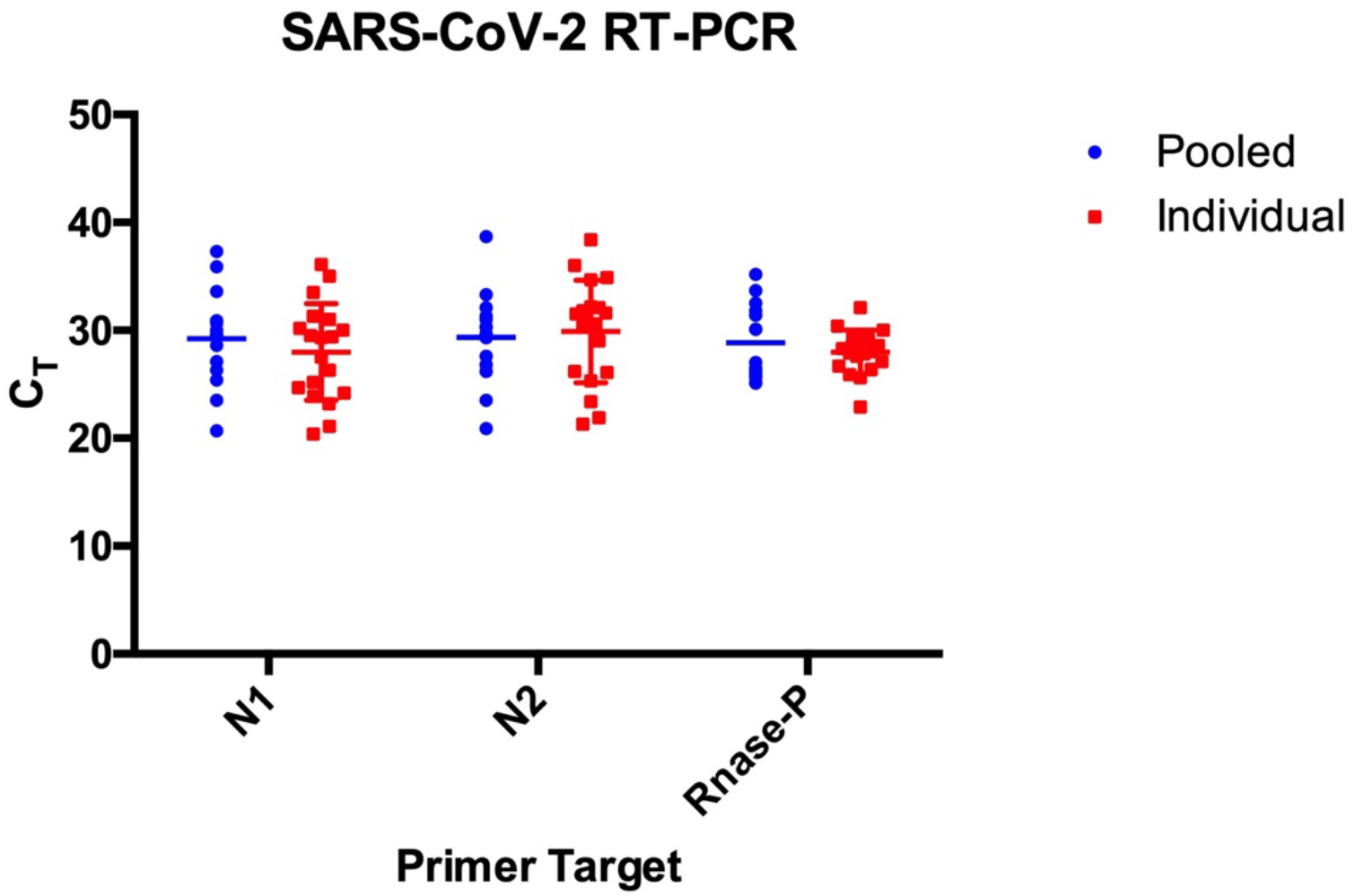
Comparison of mean C_T_ value between positive pooled and individually tested samples. Data are represented as the mean ± standard error the mean.

To determine if a pooling approach is feasible with rapid, moderate complexity tests, we tested the 15 SARS-CoV-2 positive pools and 15 of the SARS-CoV-2 negative pools using the recently released BioFire COVID-19 Test. The BioFire COVID-19 test is a nested multiplexed RT-PCR test that automates all aspects of nucleic acid testing including sample preparation, extraction, and PCR, and which can detect SARS-CoV-2 within a single nasopharyngeal swab specimen in under 60 minutes. As expected, there was perfect agreement between the CDC 2019-nCoV RT-PCR and BioFire COVID-19 assays (kappa=1.0). SARS-CoV-2 RNA was detected in all 15 positive pools, whereas SARS-CoV-2 RNA was not detected in all 15 of the negative pools using the BioFire COVID-19 Test (**Table 3**).

**Table 3.** CDC 2019-nCoV RT-PCR and BioFire COVID-19 comparator analysis

|  |  | BioFire COVID-19 Test |  |
| --- | --- | --- | --- |
|  |  | Positive | Negative |
| CDC 2019-nCoV RT-PCR | Positive | 15 | 0 |
|  | Negative | 0 | 15 |

## Discussion

We found that a single NP swab specimen containing SARS-CoV-2 RNA can be consistently detected in a pool of 10 samples. Our data shows an estimated false negative rate of approximately 7% (1 out of 15), although this pool was inconclusive (the N2 primer failed to amplify) and was treated as a positive pool. Unlike other pooling strategies that pool purified RNA extracts^12,13^, our method utilized pooling clinical specimens prior to RNA extraction, which removes the extraction bottleneck and allows running an endogenous internal control to monitor extraction quality.

A linear increase in threshold cycle is expected as specimens are pooled, however, we did not observe a significant change in CT values for either primer pair in our pooled samples. Given that PCR efficiency of each primer pair can differ, any inconclusive result for a pool should be treated as positive and individually tested. Case in point, the only inconclusive result in our study was found in pool 41, where the N2 target failed to amplify. This pool contained 2 positive specimens and one inconclusive specimen. This suggests there may have been PCR inhibitors present in the individual sample that carried over to the pooled specimen resulting in an inconclusive result. Both positive specimens in pool 41 had relatively high CT values. In our lab, specimens with high CT values are commonly observed in convalescent patients 14-30 days after symptomatic infection, and do risk escaping detection when combined in larger pools due to loss of sensitivity. It should also be noted that a negative pool result would not differentiate between a true negative and an inconclusive or invalid result due to improper sample collection or storage. Given the clinical performance of this and other published pooling protocols^7,12,13^, it is possible that larger pools could be used with further RT-PCR optimization to allow lower detection limits for low-concentration RNA.

Disease prevalence should also be taken into consideration when implementing a pooling strategy. Recently, Noriega and Samore used a Bayesian modeling approach to show testing throughput more than doubles when prevalence rates are ≤8%, and this occurs with optimal pool sizes between 4 and 12 samples. Conversely, as prevalence increases, they show improvements in testing throughput diminishes significantly^14^. During this surveillance period, SARS-CoV-2 prevalence was determined to be approximately 4% (19/494), which is ideal for pool sizes of 10. In this study, we found that 30% (15/50) pools were positive for SARS-CoV-2, which equates to 200 individual extractions and RT-PCR reactions (50 pools and 150 individuals), representing a 60% savings in extractions and RT-PCR reactions, which is significant during times of surge testing and in limited-resource situations.

Recently, numerous rapid molecular diagnostic platforms have received an Emergency Use Authorization from the FDA. These include low to moderate complexity assays from BioFire, Cepheid, and Abbott that can detect SARS-CoV-2 in approximately 1 hour^15,16^. Using the recently released BioFire COVID-19 Test, we found that this platform could reliably detect a single positive sample in pools of up to 10 specimens, with equal rates of detection as the CDC COVID-19 RT-PCR assay. This is not surprising given the published limits of detection for the CDC COVID-19 RT-PCR and BioFire COVID-19 test are in the range of 10^2^ RNA copies/ml. These results support the use of rapid molecular diagnostic platforms for routine disease surveillance of critical working groups such as healthcare providers and military units, where large-scale quarantines can have grave consequences.

In summary, we show that a pooled-sample strategy can augment a laboratory’s testing capability and relieve extreme pressure from limited resource situations without sacrificing RT-PCR sensitivity and specificity. Importantly, a pooling strategy can reduce turnaround times for prompt identification and isolation of infected individuals to effectively curb the transmission of COVID-19 and other infectious disease outbreaks.

## Acknowledgements

We thank the Landstuhl Regional Medical Center Department of Virology staff for their assistance with specimen processing. The views expressed belong to the author(s) and do not represent the official views of the United States Government, Department of Defense or Landstuhl Regional Medical Center.

